# Longitudinal and large-scale monitoring of transcriptome and RBP-RNA interactome in living cells by engineered protein nanocages

**DOI:** 10.1101/2025.03.13.642977

**Authors:** Lu-Feng Hu, Gang Xie, Yi-Xia Wu, Yu-Xuan Li, Zi-Li Wan, Li-Mi, Jia-Zhen Wang, Yangming Wang

**Affiliations:** State Key Laboratory of Gene Function and Modulation Research, Institute of Molecular Medicine, College of Future Technology, Peking University, Beijing, China; PKU-THU-NIBS Joint Graduate Program, Academy for Advanced Interdisciplinary Studies, Peking University, Beijing, China; Beijing Advanced Center of RNA Biology (BEACON), Peking University, Beijing, China; Research Center for Industries of the Future, School of Life Sciences, Westlake University, Hangzhou, Zhejiang, China

## Abstract

Nondestructive sequencing of RNA from live cells is essential for monitoring and understanding dynamic biological processes. However, most existing RNA sequencing methods rely on cell lysis or fixation, limiting their applicability for longitudinal studies. Here, we introduce POND-seq (<u>P</u>rotein nanocage-emp<u>O</u>wered <u>N</u>on-<u>D</u>estructive <u>seq</u>uencing), a novel approach that employs secretory protein nanocages fused with RNA-binding proteins (RBPs) to capture the RBP-RNA interactome and transcriptome in live cells. POND-seq reliably identifies RNA targets of canonical RBPs across multiple cell types. By fusing poly(A)-binding protein (PABPC1) to the nanocage, we demonstrate that POND-seq can monitor transcriptomic changes in response to signaling stimuli and selectively capture cell-type-specific transcriptomes from mixed populations. Additionally, POND-seq facilitates the dissection of RNA-binding domains and key amino acid residues critical for RBP-RNA interactions. We further highlight its utility in large-scale screening, offering compelling evidence for the pathogenicity of FMR1 variants. POND-seq represents a transformative advancement in RNA biology, cell biology and precision medicine, enabling unprecedented insights into cellular dynamics and disease mechanisms.

## Main

High throughput RNA sequencing (RNA-seq) is a revolutionary technology for characterizing cellular states and mechanistic investigations in biomedical sciences^1,2^. However, conventional RNA-seq methods require cell lysis, making them inherently destructive and limiting their application to single snapshots in time. This prevents longitudinal studies on the same population of cells, restricting our understanding of dynamic processes such as transcriptional reprogramming, stress responses, and differentiation on the same biological samples.

Similarly, non-destructive investigation of RNA-binding protein (RBP)-RNA interactions — crucial for post-transcriptional regulation — remains technically challenging. RBPs play essential roles in RNA fate determination, and mutations in RBPs are increasingly recognized as drivers of various diseases, including cancer and neurogenetic disorders^3–5^. Understanding how RBP mutations alter RBP-RNA interactomes is key to deciphering disease mechanisms and identifying therapeutic targets. However, current methods^6,7^ for characterizing RBP-RNA interactome require labor-intensive workflows involving one or more steps of cell fixation, crosslinking, immunoprecipitation, and lysis, making them difficult to scale for high-throughput applications, for example, studying large numbers of mutations in clinical samples. Furthermore, these methods cannot provide dynamic interactome data from live cells, leaving a critical gap in our ability to track RBP functions in the same biological samples.

To address these limitations, we developed POND-seq, a <u>p</u>rotein nanocage emp<u>o</u>wered <u>n</u>on-<u>d</u>estructive RNA <u>seq</u>uencing platform that allows transcriptome analysis and RBP-RNA interactome profiling in live cells (Fig. 1a, b). By fusing RBPs with designed protein nanocage protomer EPN24^8^, POND-seq facilitates the packaging and secretion of RBP-bound RNAs into the extracellular medium, eliminating the need for cell lysis. Benchmarking against cross-linking and immunoprecipitation (CLIP) based methods, we demonstrate that POND-seq reliably identifies RBP-interacting targets across 9 different RBPs and multiple cell lines. Furthermore, by fusing EPN24 with poly A binding protein C (PABPC1)^9^, POND-seq accurately captures the transcriptome of cells, enabling longitudinal sampling of the same cell population across multiple timepoints. Notably, we showcase POND-seq’s utility in parallel screening of 77 variants of fragile X messenger ribonucleoprotein 1 (FMR1), revealing their distinct effects on RNA interactions and identifying a putative new disease-causing mutation for fragile X syndrome^10^. Altogether, with its non-destructive and scalable design, POND-seq paves the way for broad biomedical applications, such as investigating RNA biology, monitoring cell fate transitions, and elucidating disease mechanisms.

**Fig. 1:**
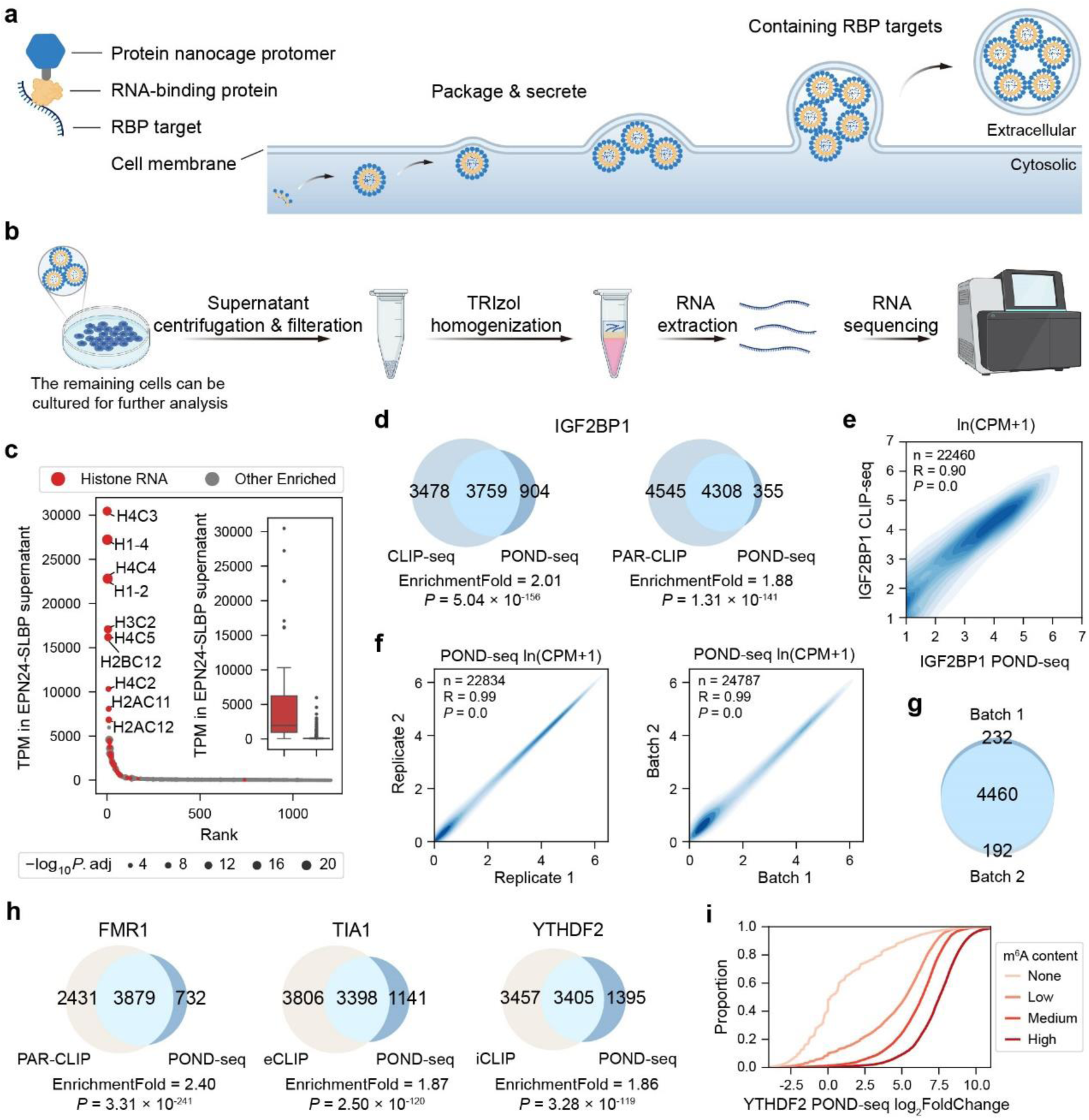
Developing and optimizing POND-seq for profiling RBP targets in living cells. **a,** Design of RBP fused protein nanocage. The protein nanocage protomer with a Myc tag (gray), fused to the N-terminal of RNA-binding protein (RBP), packs the RBP-binding RNAs into nanocages that are then secreted from cells. **b,** Experimental workflow of POND-seq. After transfection, the supernatant is collected, followed by centrifugation and filtration. RNA is then extracted with TRIzol reagent for sequencing and subsequent analysis. **c,** Rank plot with an inset box plot showing the distribution of transcripts per million (TPM) for RNAs enriched in the cell supernatant containing EPN24-SLBP nanocages. Histone RNAs are highlighted in red. The dot size represents statistical significance (EPN24-SLBP versus EPN24-GFP, adjusted *P*-value, −log10). For box plot, lower and upper hinges represent the first and third quartiles, the center line represents the median, and whiskers represent ±1.5× the interquartile range. **d,** Venn diagrams showing the overlap of targets determined by POND-seq and other methods for IGF2BP1 in HEK293T cells. **e,** Pearson’s correlations of ln(CPM+1) between IGF2BP1 POND-seq and CLIP-seq in HEK293T cells. **f,** Pearson’s correlations of ln(CPM+1) between replicates (left) and batches (right) of IGF2BP1-POND. **g,** Venn diagram showing IGF2BP1 targets identified in two independent batches of POND-seq performed about one month apart. **h,** Venn diagrams showing the overlap of targets determined by POND-seq and other methods for FMR1 (left), TIA1 (middle), and YTHDF2 (right) in HEK293T cells. **i,** Cumulative plot showing the distribution of log_2_foldchange for transcripts in EPN24-YTHDF2 versus EPN24 HEK293T cells, which were divided into 4 groups based on m^6^A content measured by GLORI (None *n* = 30,987, Low *n* = 5,085, Medium *n* = 5,085, High *n* = 5,085). The *P*-values in (**c**) were determined by Wald test in DESeq2, in (**d**) and (**h**) by two-tailed Fisher’s exact test. The correlation coefficient R and *P*-values in (**e**) and (**f**) were determined by two-tailed Pearson’s correlation test. See also Supplementary Fig.1.

## Results

### Development of POND-seq for nondestructive identification of RBP targets in live cells

We tested five different protein nanoparticles in HEK293T cells to identify those that could selectively recover RNA targets from the cell supernatant. These protein nanoparticles^8,11^ included the capsid protein of HIV-1 (HIV-gag) and its derivative HIV-gagZIP, the retrovirus-like protein mPEG10, hPEG10 and an engineered protomer EPN24, which can self-assemble into protein nanocages and secret from mammalian cells. We chose SLBP, a RBP known to primarily bind the hairpin at the end of histone mRNA^12^, to test these nanoparticles. We made constructs fusing nanoparticle proteins with SLBP, or GFP as control, and transfected HEK293T cells. We collected cell supernatant at 48 hours after transfection, extracted RNA, and performed RT-qPCR experiments. The results show that EPN24-SLBP exhibited the highest efficiency in enriching histone RNA targets of SLBP, followed by HIVgagZIP-SLBP, whereas other protein nanoparticles demonstrated significantly lower efficiency (Supplementary Fig. 1a). RNA-seq confirmed that EPN24-SLBP could package histone RNA targets into cell supernatant in a highly selective manner (Fig.1c).

Encouraged by above results, we applied POND-seq for IGF2BP1, a canonical multi-domain RBP known for its critical roles in embryogenesis and carcinogenesis^13^, in HEK293T cells. Comparison with CLIP-seq^14^ and PAR-CLIP^15^ results from the same cell line demonstrated that POND-seq can faithfully recover the RNA targets of IGF2BP1 (Fig.1d). Notably, the abundance of RNAs detected by POND-seq showed a strong correlation with the read counts of the same genes identified by CLIP-seq (Fig. 1e). Here we found that a significant portion of reads are mapped to rRNAs (Supplementary Fig. 1b). Although rRNA depletion was not strictly necessary for identifying target RNAs, it increased the proportion of informative reads, thereby reducing sequencing costs (Supplementary Fig. 1b-d). As a result, an rRNA depletion step was included in all subsequent experiments in this study. In addition, POND-seq demonstrated high reproducibility in RBP target enrichment across replicates (samples from different wells transfected at the same time) and batches (samples from different batches transfected at different times), underscoring its robustness (Fig. 1f, g).

Next, we tested three other parameters that may impact the effectiveness of POND-seq. First, using EPN24-GFP or EPN24 as controls showed minimal impact on RNA target identification (Supplementary Fig.1e, f). Second, the number of identified targets were generally increased with longer duration of nanoparticle production (Supplementary Fig. 1g), however, the performance of POND-seq with 8 hours of accumulation was only slightly worse than 48 hours. Finally, we expressed EPN24-IGF2BP1 fusion protein at different levels and found that EPN24-IGF2BP1 at the level as low as ~10% of the endogenous IGF2BP1 was effective to capture IGF2BP1 targets (Supplementary Fig. 1h, i).

To evaluate the versatility of POND-seq across different RBPs, we applied it to enrich RNA targets of several well-characterized RBPs, including FMR1^16^, TIA1^17^, G3BP1^18^, PUM2^19^, and YTHDF2^20^ (Fig. 1h and Supplementary Fig. 1j). The results obtained from POND-seq showed high concordance with those from classical methods^16–20^ such as CLIP and eCLIP (Fig. 1h and Supplementary Fig.1j). Moreover, consistent with the roles of YTHDF2^21^, IGF2BP1^22^, and FMR1^23^ as m^6^A readers, transcripts with higher m^6^A levels determined by GLORI^24^ were more strongly enriched by POND-seq for these RBPs (Fig.1i and Supplementary Fig.1k). Furthermore, we captured RNA targets of LIN28A in both mouse and human embryonic stem cells (ESCs), finding strong agreement with previous CLIP-seq results^14,25^ (Supplementary Fig. 1l). These findings demonstrate the broad applicability of POND-seq across diverse RBPs and cell types. Collectively, our results highlight POND-seq’s robustness and versatility in investigating RBP-RNA interactions across various experimental contexts.

### Development of POND-seq for nondestructive monitoring of the transcriptome in live cells

Our next objective was to engineer a protein nanocage capable of non-specifically packaging cytoplasmic transcripts, enabling comprehensive analysis of the transcriptome in live cells. To achieve this, we tested two non-specific single-stranded RNA-binding proteins, ORF5 (open reading frame 5)^26^ and hBD3 (β-defensin 3)^27^, alongside PABPC1^9^, which predominantly binds to the 3’ poly(A) tail of eukaryotic mRNAs. Among these, EPN24 fused with PABPC1 showed the best performance in capturing transcriptome information in HEK293T cells (Fig. 2a, b and Supplementary Fig. 2a). Notably, ectopic expression of EPN24-PABPC1 had minimal impact on the transcriptome profile or proliferation rate of HEK293T cells (Supplementary Fig. 2a, b). Furthermore, even when EPN24-PABPC1 was expressed at levels as low as ~20% of endogenous PABPC1, its ability to capture the transcriptome information remained largely unaffected compared to higher levels of EPN24-PABPC1 expression (Supplementary Fig. 2c, d).

**Fig. 2:**
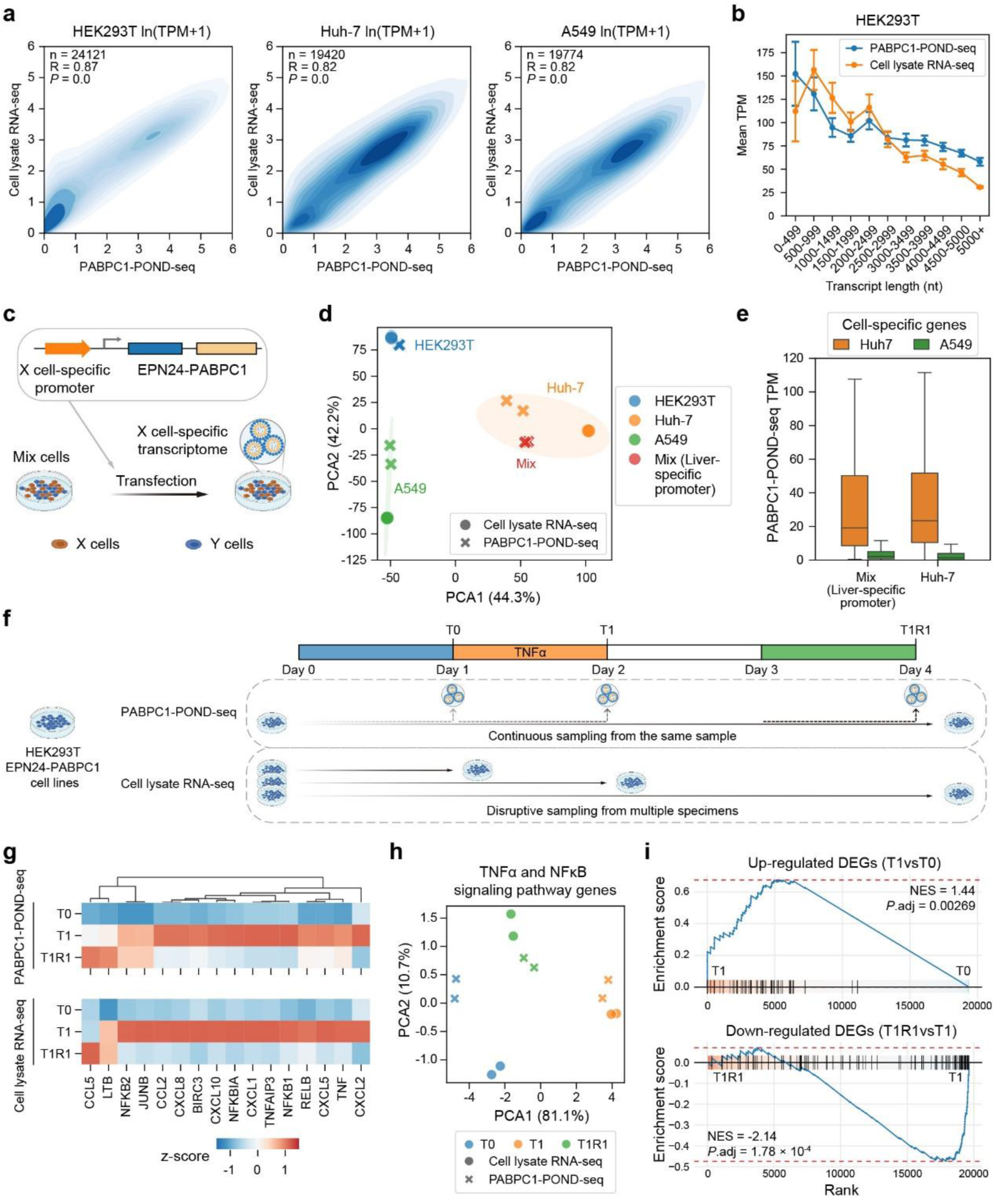
Profiling transcriptome in living cells with PABPC1 fused protein nanocage. **a,** Pearson’s correlations between cell lysate RNA-seq and PABPC1-POND-seq in HEK293T (left), Huh-7 (middle), and A549 (right) cells. Data shown are averages of two biological replicates. **b,** Line plots illustrating the mean transcripts per million (TPM) of expressed coding transcripts (TPM >1) in HEK293T cells, as determined by cell lysate RNA-seq and PABPC-POND-seq, across bins of transcript lengths. Each data point represents the mean ±SD TPM value for transcripts within the corresponding length bin. Shown are averages of two biological replicates. **c,** Schematic illustrating the determination of cell type-specific transcriptome from a mixed cell population through the cell type-specific expression of EPN24-PABPC1 by cell type-specific promoters. **d,** PCA analysis of gene expression by PABPC1-POND-seq and cell lysate RNA-seq in HEK293T, Huh-7, and A549 cells. PABPC1-POND-seq by liver-specific promoter to capture Huh-7-specific transcriptome in mixture of cells is labeled as red. **e,** Box plots showing TPM for Huh-7 and A549 cell-specific gene sets in mixed cell PABPC1-POND-seq using a liver-specific promoter and in Huh-7 PABPC1-POND-seq using a constitutive CAGGS promoter. Genes with TPM > 10 in Huh-7 cells and TPM < 1 in A549 cells are defined as Huh-7 cell-specific, and vice versa for A549 cell-specific genes. Lower and upper hinges represent the first and third quartiles, the center line represents the median, and whiskers represent ±1.5× the interquartile range. **f,** Workflow depicting TNFα treatment of HEK293T cells, followed by cell lysate RNA-seq and PABPC1-POND-seq for transcriptome analysis. **g,** Heatmaps displaying changes in core TNF signaling pathway genes during TNFα treatment and recovery, as determined by PABPC1-POND-seq (top) and cell lysate RNA-seq (bottom). Data shown are averages of two biological replicates. **h,** PCA analysis of gene expression for TNF and NFκB signaling pathways during TNFα treatment and recovery in HEK293T cells, as determined by PABPC1-POND-seq and cell lysate RNA-seq. **i,** Enrichment profiles showing DEGs identified by cell lysate RNA-seq up-regulated and down-regulated significantly in PABPC1-POND-seq after TNFα treatment and recovery. The correlation coefficient R and *P*-values in (**a**) were determined by two-tailed Pearson’s correlation test. See also Supplementary Fig.2.

We also tested EPN24-PABPC1 in A549 and Huh-7 cells and observed that PABPC1-POND-seq performed much worse in these cell types compared to HEK293T cells (Supplementary Fig. 2e). Detailed analysis of sequencing read distributions revealed substantial genomic DNA contamination in the samples from these cells (Supplementary Fig. 2f). We hypothesized that this issue stemmed from the difficulty of transfecting A549 and Huh-7 cells, which resulted in lower RNA content in the supernatant and amplified the impact of genomic DNA contamination. To address this, we introduced DNase I treatment to remove genomic DNA prior to RNA library preparation. After this adjustment, PABPC1-POND-seq successfully captured the transcriptomes of both A549 and Huh-7 cells, achieving performance comparable to that observed in HEK293T cells (Fig. 2a and Supplementary Fig. 2g).

Obtaining transcriptome information from specific cell types without requiring isolation steps from mixed cell populations is crucial for many biological studies. To assess whether POND-seq can selectively capture the transcriptome of target cells within a mixed cell population, we transfected EPN24-PABPC1 driven by a liver-specific ApoE/hAAT synthetic promoter^28^ into Huh-7 and A549 cell mixtures and conducted POND-seq (Fig. 2c). Principal component analysis (PCA) and marker gene profiling confirmed that POND-seq selectively captured the transcriptome specific to Huh-7 cells within the Huh-7/A549 mixtures (Fig. 2d, e). These findings highlight the promising potential of POND-seq for studying the transcriptomes of target cells within heterogeneous cell populations.

One of the key advantages for POND-seq is its ability to extract transcriptome information without damaging cells, enabling longitudinal monitoring of the transcriptome within the same samples. To demonstrate this capability, we treated HEK293T cells expressing EPN24-PABPC1 with TNFα^29^ for one day, followed by a two-day recovery period (Fig. 2f). Cell lysates and supernatants were collected daily for RNA-seq and POND-seq analysis (Fig. 2f). POND-seq successfully captured gene expression changes related to the TNF signaling pathway corresponding to TNFα treatment and subsequent recovery (Fig. 2g and Supplementary Fig. 2h). PCA analysis revealed a strong correlation between POND-seq and RNA-seq data, demonstrating that POND-seq effectively monitors transcriptome dynamics during cell state changes upon TNFα stimulation and withdrawal (Fig. 2h). Notably, differentially expressed genes identified by RNA-seq were significantly enriched in the POND-seq dataset (Fig. 2i and Supplementary Fig.2i). These results highlight the robustness of POND-seq for capturing dynamic transcriptome changes in living cells over an extended period, showcasing its potential for studying cellular responses to external stimuli and monitoring temporal changes in gene expression.

### Functionally characterizing RBP domains and residues by POND-seq

We next explored whether POND-seq could be utilized to quantitatively dissect the RNA-binding mechanism of RBPs. Previous studies have shown that the KH domains in IGF2BP1 play critical roles in its RNA-binding ability, particularly KH3-4 di-domain^30^. To quantitatively compare RNA content captured by POND-seq for IGF2BP1, IGF2BP1^ΔKH1-2^, and IGF2BP1^ΔKH3-4^, we first normalized RNA species by quantifying protein levels of EPN24 fusion proteins in supernatant with western blotting (Supplementary Fig. 3a). The results indicated that deletion of the KH1-2 or KH3-4 domains caused a significant global reduction in IGF2BP1-bound RNAs (Fig. 3a, Supplementary Fig. 3b). Notably, some RNA targets exhibited a domain-dependent regulation, with certain targets being more affected by the deletion of KH1-2 (KH1-2– dominant) and others by KH3-4 (KH3-4–dominant) (Fig. 3a). Interestingly, the KH3-4–dominant genes had higher m^6^A content and showed greater downregulation upon IGF2BP1 knockdown (Fig. 3b), underscoring the pivotal role of the KH3-4 domains in the m6A reader function of IGF2BP1^22^. GO enrichment analysis further revealed KH3-4–dominant genes are involved in fundamental biological processes, including RNA splicing and chromosome segregation (Fig. 3c).

**Fig. 3:**
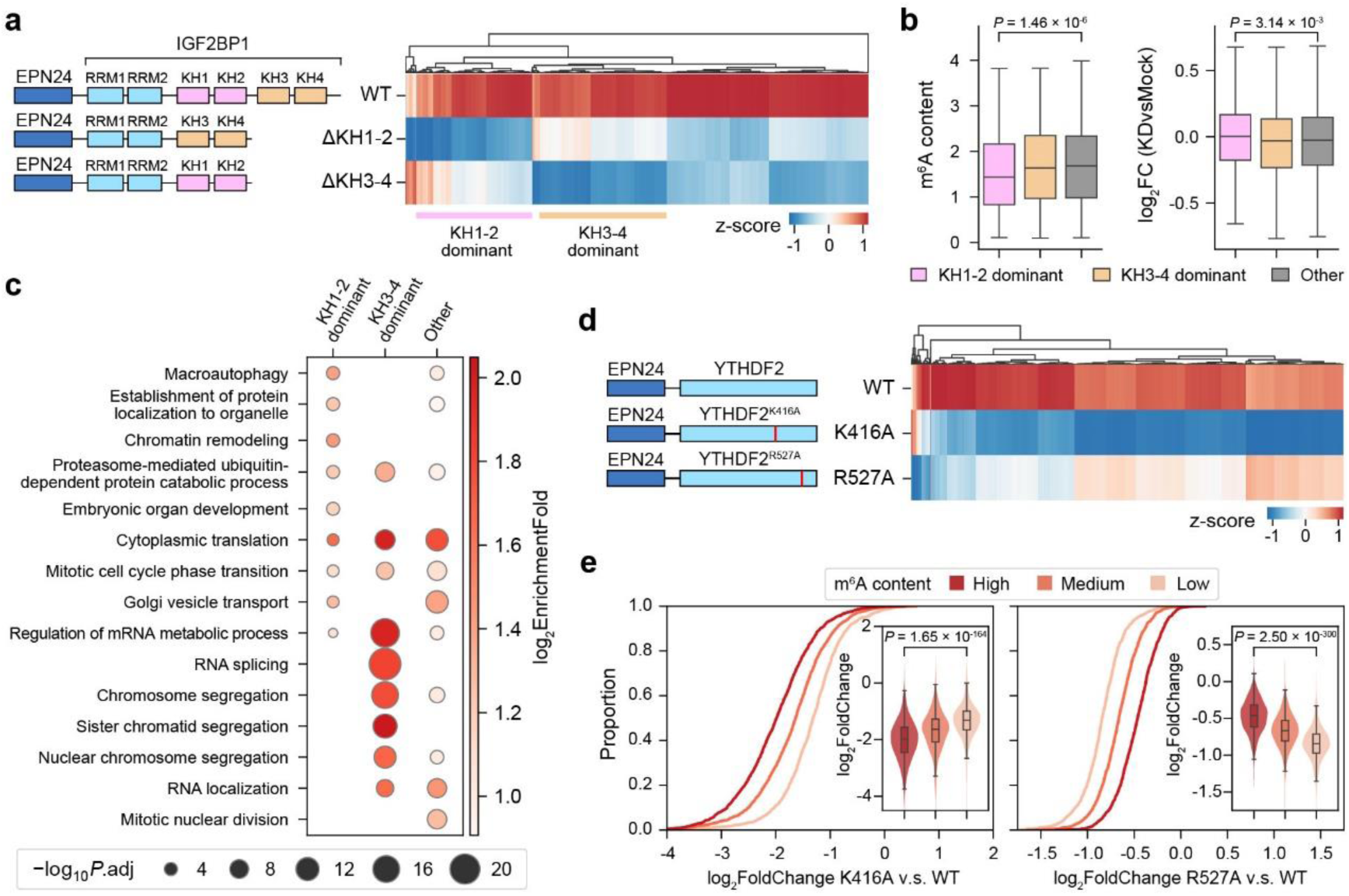
Dissecting the role of critical domains and residues in RBP-RNA interactions by POND-seq. **a,** Schematic representation (left) of the three IGF2BP1 constructs fused with EPN24: wild-type (WT), KH1-2 domain deletion (ΔKH1-2), and KH3-4 domain deletion (ΔKH3-4). The heatmap (right) shows the enrichment of WT IGF2BP1 targets across these constructs in POND-seq. Targets with a significant decrease in binding upon deletion of either the KH1-2 or KH3-4 domains are classified as KH1-2– dominant or KH3-4–dominant, respectively. **b,** Box plots showing the distribution of m^6^A content (left), and log2foldchange (si-IGF2BP1 versus mock, right) for KH1-2–dominant, KH3-4–dominant, and other genes. The lower and upper hinges represent the first and third quartiles, the center line represents the median, and whiskers represent ±1.5× the interquartile range. **c,** GO enrichment analysis of KH1-2–dominant, KH3-4–dominant, and other target genes. Top 5 GO terms for each set were selected to visualize. The dot size represents statistical significance (adjusted *P*-value, −log_10_) and the color represents enrichment fold (log_2_). **d,** Schematic representation(left) of three YTHDF2 constructs fused with EPN24: WT, K416A, and R527A. The heatmap (right) shows the enrichment of WT YTHDF2 targets across these constructs in POND-seq. **e,** Cumulative and box-violin (inset) plots showing the distribution of log_2_foldchange for YTHDF2 bound transcripts in EPN24-YTHDF2^K416A^ (left) and EPN24-YTHDF2^R527A^ (right) versus EPN24-YTHDF2 (WT) HEK293T cells. Transcripts were divided into 3 groups based on m^6^A content measured by GLORI (Low, *n* = 1,563; Medium, *n* = 1,564; High, *n* = 1,563). The *P*-values in (**b**) and (**e**) were determined by two-tailed Wilcoxon rank-sum test. See also Supplementary Fig.3.

Next, we focused on YTHDF2 to pinpoint the essential amino acid residues responsible for its RNA-binding activity. Previous studies have established that K416 and R527 residues are critical for the RNA-binding function of YTHDF2, with mutations at either site significantly impairing this ability^31^. To investigate further, we conducted POND-seq on YTHDF2 carrying K416A or R527A substitutions. While neither mutation affected the levels of EPN24-YTHDF2 protein in the supernatant (Supplementary Fig. 3c), both YTHDF2^K416A^ and YTHDF2^R527A^ exhibited reduced RNA enrichment compared to wild-type YTHDF2 (Fig. 3d and Supplementary Fig. 3d). Consistent with prior findings^31^, we observed that both mutations diminished YTHDF2 binding to m^6^A-modified RNA transcripts (Fig. 3e). Notably, the K416A mutation more severely disrupted YTHDF2’s interaction with highly m^6^A-modified RNA transcripts, whereas the R527A mutation had a stronger impact on low m^6^A-modified RNA transcripts (Fig. 3e and Supplementary Fig. 3e, f). Together with the analysis of IGF2BP1 domain deletions, these findings underscore POND-seq’s utility as a powerful tool for dissecting the molecular determinants of RBP-RNA interactions and their functional implications. By enabling quantitative comparisons of binding activity across RBP variants, POND-seq provides unique insights into the mechanisms by which specific amino acid residues or structural domains contribute to RNA binding specificity and regulation.

### Adaptation of POND-seq for large-scale screening of pathogenic mutations in FMR1

Large-scale sequencing projects generate vast amounts of genetic data, offering an invaluable resource for identifying previously unknown mutations in RNA-binding proteins (RBPs) that may contribute to disease. However, pinpointing disease-causing mutations and understanding their underlying mechanisms remain significant challenges, primarily due to the cumbersome and time-consuming procedures of traditional methods such as CLIP-seq and eCLIP. These techniques require complex sample preparation and labor-intensive analyses, limiting their applicability in large-scale mutation screenings. In contrast, POND-seq offers a more streamlined and scalable approach. Here, we explored whether POND-seq can be automated for high-throughput screening, enabling more efficient identification of disease-causing mutations and providing insights into the molecular mechanisms of RBP-related diseases. As a proof of concept, we focused on FMR1 (Fig. 4a), a disease-associated RNA-binding protein known for its central role in Fragile X syndrome (FXS), the most common monogenetic cause of intellectual disability^10,32^. While the majority of FXS cases are linked to abnormal CGG repeat expansions in the 5’-untranslated region of FMR1, a subset of patients with normal CGG repeat lengths still exhibit FXS-like symptoms^33–35^. Sequencing studies of these patients have identified missense mutations^36–39^, highlighting the need to investigate the relationship between these non-CGG expansion mutations and the clinical manifestations of FXS.

**Fig. 4:**
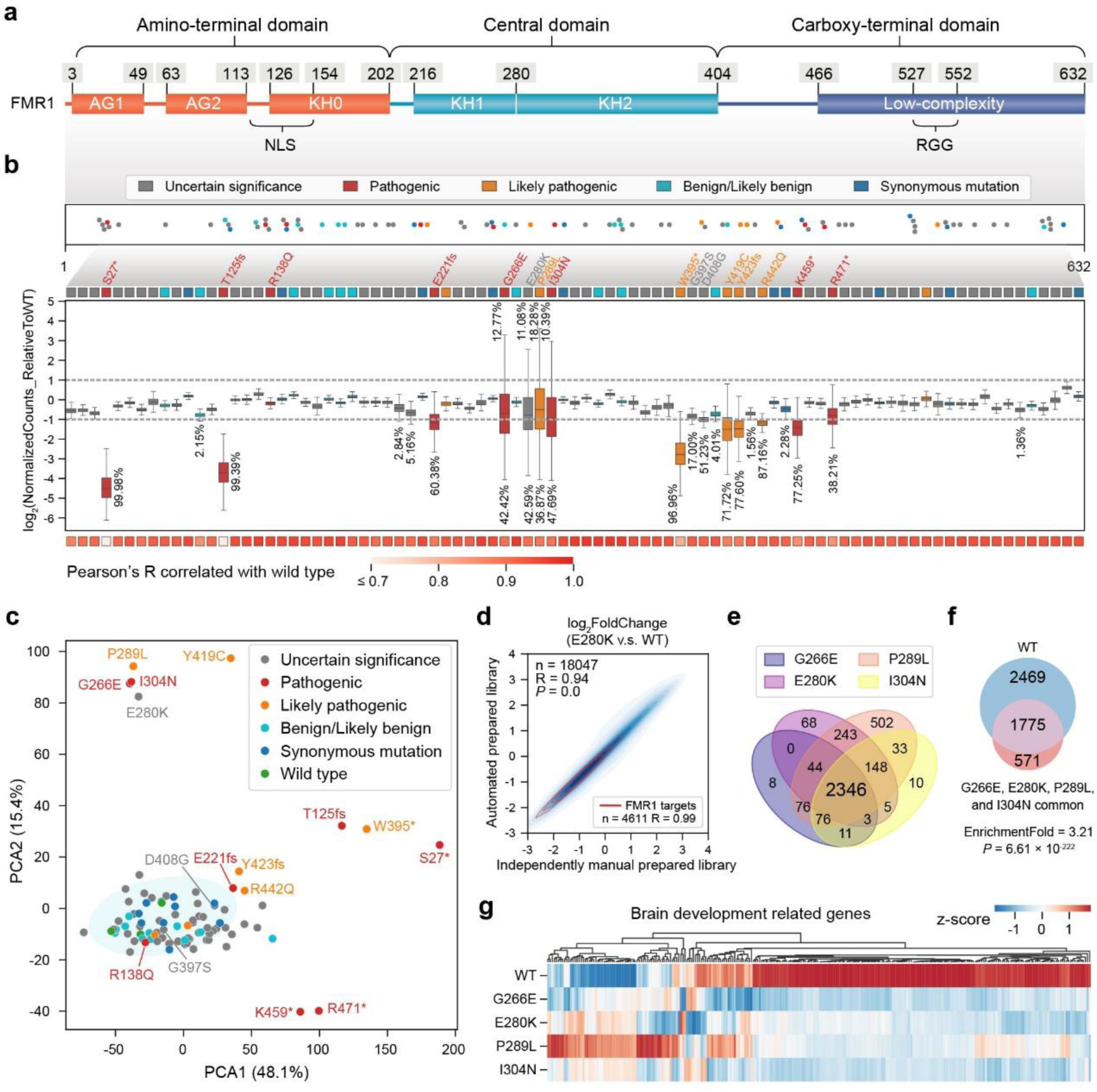
Large scale screening by POND-seq identifies putative new pathogenic mutations in FMR1. **a,** Molecular and functional domains of FMR1. Three major domains are color-coded, and numbers indicate position within the amino acid sequence. **b,** Scatter plots (top) depicting FMR1 mutation sites analyzed by POND-seq, with the mutation positions corresponding to locations shown in (**a**). Box plots (middle) illustrating the distribution of normalized counts for all genes determined by POND-seq of FMR1 variants relative to wild-type FMR1 POND-seq. The fill color indicates the pathogenicity of the mutation sites. The lower and upper hinges represent the first and third quartiles, the center line indicates the median, and whiskers extend to ±1.5× the interquartile range. A heatmap (bottom) displaying the Pearson’s correlation coefficients between interactomes of variant FMR1 and wild-type FMR1. **c,** PCA analysis for POND-seq data of WT and variant FMR1. **d,** Pearson’s correlation analysis of log2foldchange in transcript levels between FMR1 E280K and WT FMR1 POND-seq, comparing libraries prepared using automated versus manual methods. **e,** Venn diagrams illustrating the overlap of RNA targets bound by FMR1 variants (G266E, E280K, P289L, and I304N) as identified by POND-seq. **f,** Venn diagrams illustrating the overlap of common RNA targets bound by all four variants (G266E, E280K, P289L, and I304N) and wild-type FMR1 as identified by POND-seq. **g,** Heatmap showing enrichment of the brain development related gene transcripts in POND-seq of WT, G266E, E280K, P289L, and I304N variants. The correlation coefficient R and *P*-values in (**d**) were determined by two-tailed Pearson’s correlation test. The *P*-values in (**f**) were determined by two-tailed Fisher’s exact test. See also Supplementary Fig.4.

EPN24-FMR1 mutants were cloned and expressed in HEK293T cells, followed by supernatant collection and RNA extraction. In total, 68 missense mutations, 4 nonsense mutations, 3 frameshift mutations, and 2 microdeletion mutations listed in the NCBI-ClinVar database^36^ were generated for analysis. Additionally, 10 synonymous mutations were included to evaluate the false positive rate of the POND-seq screening assay. Leveraging an automated library preparation workstation, POND-seq libraries for supernatant RNA of 182 samples (87 mutations, 3 wild type, 1 EPN24 control, and 2 replicates for each construct) were completed in just 12 hours, a significant improvement over conventional CLIP-based methods, which would require months to process the same number of samples.

Sequencing results showed that seven of the eight pathogenic mutations significantly impaired RNA binding, with 38.21% to 99.98% of targets reduced by more than twofold (Fig. 4b). The exception was R138Q, which showed only a modest decrease. This observation aligns with previous studies indicating that FMR1-R138Q retains RNA-binding activity and translational regulation but causes FXS phenotypes through an unclear mechanism^38^. Of the seven mutations annotated as likely pathogenic, five showed substantial reductions in RNA-binding ability (36.87% to 96.96% of targets reduced by more than twofold), while two mutations (V225I and Q541_G545del) did not exhibit significant changes in RNA binding (Fig. 4b). Importantly, all 11 benign/likely benign mutations, as well as the 10 synonymous mutations, had minimal effects on the FMR1-RNA interactome (Fig. 4b). These results highlight the reliability of POND-seq in characterizing disease-associated mutations.

Among the mutations of uncertain significance, three variants-E280K, G397S, and D408G, showed impaired RNA-binding abilities, with 42.59%, 17.00% and 51.23% of their targets, respectively, showing a reduction of more than twofold in abundance (Fig. 4b). However, PCA analysis revealed that only E280K displayed distinct RNA-binding profiles compared to wild-type FMR1 (Fig. 4c). Independent experiments were conducted to validate the effect of E280K (Fig. 4d).

E280K is a rare variant previously identified in a patient exhibiting FXS-like symptoms^36^. While E280 is predicted to be highly conserved across vertebrates and the E280K mutation is predicted to be damaging, its pathogenicity remains uncertain due to limited experimental evidence. Here, we provide new data supporting the potential pathogenicity of E280K mutation. Notably, E280 resides within KH1-KH2 domain (Fig. 4a and Supplementary Fig. 4a), a critical region for RNA binding^40^. In this domain, three adjacent mutations — G266E, P289L, and I304N — have been classified as pathogenic or likely pathogenic^36^. Interestingly, E280K exhibited similar effects on the RBP-RNA interactome as these three pathogenic mutations, based on PCA analysis (Fig. 4c).

The observed reduction in RNA enrichment was not due to secretion defects of the nanoparticles, as the protein content in the supernatant was unaffected for these mutations (Supplementary Fig. 4b, c). RT-qPCR analysis further confirmed that RNA levels of four known targets were significantly reduced in the supernatant of FMR1 harboring these four mutations compared to wild-type FMR1 (Supplementary Fig. 4d). Target analysis revealed that E280K and the three pathogenic mutations shared highly similar RNA-binding profiles (Fig. 4c, e), which differed substantially from the wild-type FMR1 profile (Fig. 4f, Supplementary Fig. 4e). Importantly, we found that the interactions between FMR1 and transcripts of brain development-related genes^41^ were similarly disrupted by E280K and the other three mutations (Fig. 4g), suggesting that E280K may be as detrimental as G266E, P289L, and I304N to processes involved in brain development.

These findings support the hypothesis that E280K contributes to FXS-related disorders through a mechanism akin to G266E, P289L, and I304N. Together, these results underscore the utility of POND-seq as a robust tool for screening RBP mutations, identifying those that impair RBP-RNA interactions, and elucidating their impact on the biological roles of RBPs.

## Discussion

In this study, we developed POND-seq, a scalable and non-destructive RNA-seq platform that utilizes engineered protein nanocages. We demonstrated its effectiveness in reliably capturing RBP-RNA interactome for nine different RBPs and in five different cell lines. By fusing PABPC1 to nanocages, POND-seq can also be customized for monitoring transcriptome dynamics in living cells. For its non-destructive nature, POND-seq allows continuous tracking of both transcriptome and RBP-RNA interactome over time, providing a unique ability to monitor the evolution of gene regulation in the same samples. The simplicity of the method makes POND-seq highly scalable and amenable to automation, facilitating high-throughput analysis of a large number of treatments, multiplexed RBP arrays, and RBP variants. By capturing key molecular mechanisms of RBP function and identifying novel pathogenic mutations, POND-seq has the potential to transform RNA biology, transcriptomics, and precision medicine, providing unprecedented insights into cellular processes in both healthy and disease contexts.

We selected EPN24 for developing POND-seq due to its effective enrichment of a select group of histone RNA targets of SLBP. While further systematic testing is required, EPN24 nanocages offer several advantages over traditional viral capsid-based virus-like particles (VLPs). Unlike VLPs, which are prone to non-specific RNA binding due to their larger size and inherent RNA-binding domains, EPN24 nanocages are smaller and less likely to bind RNA non-specifically, thereby reducing the chances of nonspecific RNA encapsulation^8^. Additionally, EPN24’s modular and customizable nature allows for precise fusion with RNA-binding proteins (RBPs), ensuring that their structural integrity is preserved^42,43^. This design enhances the specificity of RNA binding, minimizes background interference, and reduces the potential for unwanted interactions with host-cell proteins—common issues with viral VLPs^44,45^. Furthermore, by minimizing viral sequence content^8,42,43^, EPN24 nanocages mitigate immunogenic risks, making them a safer and more versatile option for potential in vivo applications.

POND-seq’s non-destructive approach offers significant advantages for studying valuable restricted cell samples, such as non-dividing, post-mitotic cells (e.g. neurons and cardiomyocytes), where traditional methods are limited due to the inability to repeatedly sample the same cells. This makes POND-seq especially valuable in studying long-term cellular dynamics in these cells. Furthermore, POND-seq is well-suited for tracking cellular transitions in complex models, such as 3D organoids^46^ or during cellular differentiation. These models often exhibit high variability between samples^47–49^, which complicates comparisons across different timepoints. By enabling serial measurements from the same biological instance, POND-seq allows for more accurate tracking of molecular changes and provides deeper insights into cellular dynamics during processes like development, differentiation, and disease progression.

POND-seq’s high-throughput capabilities make it a powerful tool for large-scale studies. Its streamlined protocol and ability to analyze multiple RBPs in parallel are particularly suited for investigating mutation effects across a broad range of disease-associated RBPs. For instance, using POND-seq, we screened 77 variants of the FMR1, identifying both previously reported mutations and novel sites that affect RBP binding. This makes POND-seq an essential tool for uncovering new pathogenic mutations and deepening our understanding of RNA-related diseases. Moreover, when combined with high-throughput drug screening, POND-seq holds the potential to identify novel therapeutics that restore the function of mutant RBPs, offering new avenues for targeted treatments^50,51^.

Our study demonstrates that PABPC1-POND-seq can selectively capture transcriptomic information from target cells within a mixed population. By expressing the EPN24 fusion protein under a cell type-specific promoter, we can efficiently isolate RNA data from distinct cell types within heterogeneous environments. This capability highlights POND-seq’s potential for obtaining gene expression data from individual cell populations within complex tissues, such as 3D organoids, thus opening new possibilities for cellular and tissue-level analysis. Looking ahead, we envision POND-seq being used in living animals, enabling longitudinal, physiological monitoring of RNA interactions in the context of disease and development. While challenges remain—such as efficiently retrieving secreted particles in vivo—the ability to apply POND-seq ex vivo may offer a promising route in more physiologically relevant settings.

While POND-seq offers many advantages, we recognize that there are some limitations for improvement in future developments. One limitation of POND-seq is its inability to directly identify RNA-binding motifs of RBPs. Since POND-seq captures full-length RNAs without enriching for the specific RNA fragments directly bound by RBPs, it cannot provide precise information about the exact binding sites on the RNA. Another limitation is the potential effect of EPN24-RBP overexpression on endogenous RNA substrates. However, this issue can be mitigated by expressing the EPN24-RBP fusion proteins at levels significantly lower than endogenous RBP levels, ensuring minimal disruption to cellular processes. Lastly, POND-seq captures integrated signals over the duration of sampling, rather than providing a snapshot at a single time point. Although we have tested relatively short time windows, the time resolution of POND-seq may not be sufficient for experiments requiring rapid, real-time monitoring of dynamic cellular changes.

In summary, POND-seq offers a revolutionary approach to RNA-seq, enabling non-destructive, longitudinal monitoring of both RBP-RNA interactome and transcriptome. With its potential for large-scale applications and its ability to study complex biological systems, we believe POND-seq will have a transformative impact across RNA biology, cell biology, and clinical research.

## Methods

### Cell culture

HEK293T, Huh-7, and A549 cells were cultured at 37°C under 5% CO_2_ in high glucose Dulbecco’s Modified Eagle’s Medium (DMEM, Hyclone) supplemented with 10% FBS (PANSera), penicillin (100 U/ml) and streptomycin (100 µg/ml). V6.5 mouse ESCs were grown on gelatin-coated dish at 37°C under 5% CO_2_, in KnockOut^TM^ DMEM (Gibco) supplemented with 15% FBS (Hyclone), nonessential amino acids, 2-mercaptoethanol, and 1,000 units/ml lymphocyte inhibitory factor. H1 Human ESCs were grown in mTeSR^TM^ (STEMCELL Technologies) on Matrigel-coated dish at 37°C under 5% CO_2_.

### Plasmid construction

EPN24-RBPs were driven by a CAGGS promoter in a piggyBac vector. Dox inducible EPN24-LIN28A and EPN24-PABPC1 were driven by a doxycycline-inducible TRE3G promoter in a piggyBac vector which harbors an rtTA trans-activator driven by a EF1α promoter. A Myc tag was inserted between EPN24 and RBPs in all EPN24-RBP constructs. The EPN24 (containing a Myc tag at C-terminal), HIVgag-ZIP, mPEG10, and hPEG10 constructs were synthesized by Azenta (Beijing). HIVgag was a gift from Antony K. Chen laboratory. The coding sequences (CDS) of RBPs were PCR amplified from HEK293T cDNA with Phanta Super-Fidelity DNA Polymerase (Vazyme). The FMR1 mutants were generated from wild-type sequence using overlap PCR-mediated site-directed mutagenesis. The EPN24 with a Myc tag and RBP CDS were assembled into vectors using 2×MultiF Seamless Assembly Mix (ABclonal). All constructs were confirmed by sanger sequencing.

### Cell transfection

For POND-seq in HEK293T, 150,000 HEK293T cells per well were seeded in a well of poly-D-lysine-coated 24-well plates. After 16-20 h, cells were transfected with 500 ng EPN24-RBPs using JetPRIME reagent (Polyplus), the medium was refreshed at 4-6 h post-transfection, and the supernatant was collected for subsequent analysis 48 h after medium change. To test POND-seq with different duration of nanoparticle production, 500 ng CAGGS-EPN24-IGF2BP1 were transfected, with medium refreshed 6 h later, and supernatant collected 8, 16, 24, 36, 48 or 72 h after medium change. To test RNA enrichment under low dosage transfection, 50 or 100 ng CAGSS-EPN24-IGF2BP1 were transfected, with medium refreshed 4 h later, and supernatant collected 48 h after medium change. To scan FMR1 mutations, 100 ng CAGSS-EPN24-FMR1 plasmids per well were transfected in 24-well plates, the medium was changed 4 h later, and the supernatant was collected 48 h after medium change.

For POND-seq in mESCs, 100,000 mESCs were seeded in gelatin-coated 12-well plates. After about 20 h, cells were transfected with 750 ng TRE3G-EPN24-Lin28A and 250 ng pBase plasmids with Lipo3000 reagent (Life Technology). The medium was refreshed 4 h later. Two days later, cells were selected and expanded in medium containing 150 μg/ml Hygromycin (Roche) with daily medium change for one week. Then 100,000 mESCs were seeded in gelatin-coated 12-well plates. About 20 h later, cells were cultured with fresh medium containing 1 μM doxycycline for two days before cell supernatant was collected for analysis. Similar procedure was conducted for POND-seq in hESCs.

For longitudinal PABPC1-POND-seq in HEK293T cells, 150,000 HEK293T cells were seeded per well in poly-D-lysine-coated 24-well plates. After about 20 h, 500 ng of TRE3G-EPN24-PABPC1 and 250 ng of pBase plasmids were transfected using JetPRIME reagent. The medium was refreshed 4 h later. Two days post-transfection, cells were selected and propagated in medium containing 150 μg/ml Hygromycin for one week. A total of 300,000 selected cells were seeded into 6-well plates. The following day, cells were cultured with fresh medium containing 1 μM doxycycline for 24 h, before the supernatant was collected. Cells were then washed three times with PBS and cultured in fresh medium with 1 μM doxycycline and 100 ng TNFα for 24 hours. The supernatant was collected again, and remaining cells were washed three times with PBS before being cultured for an additional two days with 1 μM doxycycline. During this period, cells were refreshed daily with medium containing 1 μM doxycycline, and supernatant from the last day was collected for analysis. Cells cultured in parallel under the same conditions were collected for RNA extraction and RNA-seq analysis.

For POND-seq in Huh-7 and A549, 75,000 cells were seeded in 24-well plates. After about 20 h, cells were transfected with 500 ng EPN24-PABPC1 using JetPRIME reagent. The medium was refreshed 4 h later, and the supernatant was collected for analysis 48 h after medium change. For POND-seq in Huh-7/A549 mixtures, 75,000 Huh-7 and 75,000 A549 cells were mixed and seeded in 12-well plates. After about 20 h, cells were transfected with 1 μg EPN24-PABPC1 using JetPRIME reagent. The medium was refreshed 4 h later, and the supernatant was collected for analysis 48 h after medium change.

### POND-seq

Cell supernatant was clarified by centrifugation at 3,000 g for 5 minutes, then filtrated through a cellulose acetate filter with 0.45 μm pore size (Corning). 200 μl of supernatant was mixed with 800 μl TRIzol reagent (Magen) and 1 μl diluted External RNA Controls Consortium (ERCC) synthetic spike-in RNAs (ThermoFisher). RNA was extracted and dissolved in 15-20 μl nuclease-free water. For NGS sequencing, 5 μl RNA was used for library construction with modified VAHTS Universal V6 RNA-seq Library Prep Kit for Illumina (Vazyme). Typically, 5 μl RNA was mixed with 5 μl 2×Prime buffer (Vazyme) for fragmentation at 85℃ for 6 min. For ribosomal depletion, 5 μl RNA was mixed with 5 μl 2×Prime buffer (Vazyme) and 0.7 μl FastSelect reagent (Vazyme) for fragmentation at 85℃ for 6 min. Then 8 μl fragmentated RNA was used for first and secondary strand cDNA synthesis, followed by adapter ligation and PCR amplification (16-18 cycles) according to the manufacture’s instruction manual. The DNA library was purified with VAHTS DNA Clean Beads (Vazyme) and dissolved in 20 μl nuclease-free water. The 150-nt paired-end sequencing was performed on Illumina NovaSeq 6000 (Novogene, China) or DNBSEQ-T7 platform (GenePlus, China).

For FMR1 mutation screening, library construction was completed with VNL-96P (Vazyme) automated library preparation workstation. Briefly, 4 μl RNA was mixed with 4 μl 2×Prime buffer V2 (Vazyme) and 1 μl FastSelect reagent (Vazyme) for fragmentation at 85°C for 6 min. Then 8 μl fragmentated RNA was adopted for first and secondary strand cDNA synthesis, followed by adapter ligation and PCR amplification (17 cycles) according to VAHTS Universal V10 RNA-seq Library Prep Kit for Illumina (Vazyme). The NGS sequencing was performed on DNBSEQ-T7 platform (GenePlus).

### RNA-seq

The total RNA of HEK293T, Huh-7 and A549 were extracted with TRIzol reagent (Magen) and dissolved in nuclease-free water. 500 ng total RNA was used for library construction with VAHTS Universal V6 RNA-seq Library Prep Kit for Illumina (Vazyme). The DNA library was purified with VAHTS DNA Clean Beads (Vazyme) and dissolved in 20 μl nuclease-free water. The 150-nt paired-end sequencing was performed on Illumina NovaSeq 6000 (Novogene) or DNBSEQ-T7 platform (GenePlus).

### RT-qPCR

Generally, 3 μl of supernatant RNAs or 250 ng total RNA were reverse transcribed using HiScript® III RT Super Mix (Vazyme) at 37°C for 15 minutes. The cDNA products were diluted five-fold with 20 μl nuclease-free water, and 1 μl aliquot was used for real-time qPCR with AceQ qPCR SYBR Green Master Mix (Vazyme) in 96-well plates on the StepOne Plus Real-Time PCR System (Applied Biosystems) following standard protocols.

### Western blot

Briefly, 24 μl clarified, filtrated supernatant was mixed with 6 μl 5×SDS loading buffer (Beyotime) and was boiled at 98℃ for 15-20 min in a thermal cycler. Protein electrophoresis was performed using BeyoGel™ Elite Precast PAGE Gel (8-16%, Beyotime), followed by protein transfer onto polyvinylidene fluoride (PVDF) membrane. Primary antibodies anti-Myc-tag (Beyotime, AF2867), anti-IGF2BP1(Proteintech, 22803-1-AP), and anti-PABPC1 (Proteintech, 66809-1-Ig) were diluted (1:1000) and incubated with the membrane at 4°C overnight. The anti-GAPDH (Bioss, bsm-33033M) was diluted (1:10,000) and incubated with the membrane at room temperature for 1 h. After three washes, the HRP-conjugated anti-IgG secondary antibody (Proteintech, SA00001-1) was diluted (1:10,000) and incubated with the membrane at room temperature for 1 h before treated with electrochemiluminescence (ECL) reagent (Merck Millipore, WBKLS0050). The membrane was imaged with eBLOT14 Touch Imager (e-BLOT). To assess the loading amount of protein samples, the membrane was stained with Ponceau S (Beyotime, P0022).

### POND-seq data processing and analysis

The 150-bp paired-end sequencing reads were quality-checked by FastQC (v0.11.9) and processed using Trim Galore (v0.6.7) (https://github.com/FelixKrueger/TrimGalore) to eliminate low-quality reads and trim adapter sequences. Processed reads were then aligned to the human genome (hg38) or mouse genome (mm10) combined with ERCC spike-in sequences (https://assets.thermofisher.cn/TFS-Assets/LSG/manuals/ERCC92.zip) by STAR (v2.7.10a) using the GENCODE (Human v40 or Mouse vM25) transcript annotation as transcriptome guide, with parameters: --outFilterMismatchNoverLmax 0.04. The raw counts aligned to exonic regions were quantified by Subread featureCounts (v2.0.1) with parameters: -g gene_id -t exon --primary -Q 30. For RBP POND-seq data, raw counts were normalized to counts per million (CPM) using ERCC spike-ins, and RBP targets were defined as transcripts with CPM >= 50, log2foldchange > 2, and adjusted P-value < 0.05 in RBP-POND versus control (EPN24 or EPN24-GFP) samples by DESeq2 (v1.36.0). The POND-seq data for mutated RBPs were also quantified using absolute quantification based on ERCC spike-in controls. The total CPM of wild-type RBP-POND were set to 1,000,000, and the CPM of mutated RBPs-POND were calculated relative to that of wild-type RBP. For transcriptome analysis with PABPC1-POND-seq and RNA-seq, raw counts were normalized to transcripts per million (TPM) as gene expression level. PCA was conducted by package sklearn (v1.1.1).

### Differentially expressed genes and function enrichment analysis

Differentially expressed genes (DEGs) were identified by DESeq2 (v1.36.0) in R software (v4.2.0). Gene ENSEMBL IDs were converted to Entrz IDs with R package org.Hs.eg.db (v3.15.0) or org.Mm.eg.db (v3.15.0). GO and KEGG enrichment analysis were performed using functions *enrichGO* and *enrichKEGG* in R package clusterProfiler (v4.7.1.3). GO or KEGG terms with adjusted *P*-value < 0.05 were defined as significantly enriched.

### Data visualization and statistical analysis

Package Seaborn (v0.13.2) and Matplotlib (v3.7.1) in python (v3.9.12) were used for the generation of heatmap, line plot, box-violin plot, Venn diagram, and box plot. Art schematics in Figures 1A, 1B, 2C, and 2F were created with BioRender.com. Unless otherwise specified, statistical significance between two independent groups was evaluated using a two-tailed Wilcoxon rank-sum test, performed with the scipy package (v1.10.1). To assess the significance of enrichment between RBP binding targets identified by different methods, Fisher’s exact test, also performed with the scipy package (v1.10.1), was used. The corresponding *P*-values and enrichment fold are presented below each Venn diagram.

### Data availability

All data supporting the findings of this study are available from the corresponding author on reasonable request. Sequencing data will be deposited in the Gene Expression Omnibus. Previously published data are available under accession numbers PRJNA533136 (G3BP1 PAR-CLIP in HEK293T), GSE210563 (GLORI in HEK293T), GSE230844 (YTHDF2 iCLIP in HEK293T), GSE21578 (IGF2BP1 PAR-CLIP in HEK293T), GSE39682 (FMR1 PAR-CLIP in HEK293T), GSE155729 (TIA1 eCLIP in HEK293T), GSE110519 (PUM2 CLIP-seq in HEK293T), GSE117840 (LIN28A irCLIP in mESC), GSE39872 (LIN28A CLIP-seq in hESC), and GSE21575 (IGF2BP1 knock down RNA-seq in HEK293T).

## Supporting information

Supplementary Figs. 1-4 with Figure Legends

## Acknowledgments

We thank all members in Wang laboratory for the critical reading and discussion of the manuscript. We are grateful to Dr. Antony K. Chen for providing the plasmid containing the HIVgag sequence. This study was supported by the National Key Research and Development Program of China (2021YFA1100200 to Y.W.), the National Natural Science Foundation of China (32025007 and 32130017 to Y.W.). L-F.H. was supported by Postdoctoral Foundation at Center for Life Sciences and Beijing Natural Science Foundation (5244035).

## Author contributions

L-F.H. and Y.W. conceived and designed the study. L-F.H. conducted all experiments and G.X. performed all bioinformatic analyses. L-F.H. constructed all plasmids with assistance from Y-X.L., Z-L.W., and J-Z.W. POND-seq library preparation was performed by L-F.H. and Y-X.W. L.M. drew the structural diagram of KH1-KH2 domain of FMR1. L-F.H., G.X., and Y.W. wrote the manuscript with input from other authors, and Y.W. supervised the project.

## Supplementary Figures

**Supplementary Fig.1**: Developing and optimizing POND-seq for profiling RBP targets in living cells, related to Fig.1.

**Supplementary Fig.2**: PABPC1-POND-seq captures transcriptome in a non-destructive manner, related to Fig.2.

**Supplementary Fig.3**: Functional dissection of RBP domains and residues by POND-seq, related to Fig.3.

**Supplementary Fig.4**: Large scale POND-seq screening identifies new pathogenic mutations in FMR1, related to Fig.4.

