## Supplementary Figs. 1-4 with Figure Legends for "Longitudinal and large-scale monitoring of transcriptome and RBP-RNA interactome in living cells by engineered protein nanocages"

**Affiliations:**

5 Lead author

[#]These authors contributed equally to this study.

**This file contains:**
**Supplementary Figs. 1-4 with Figure Legends**

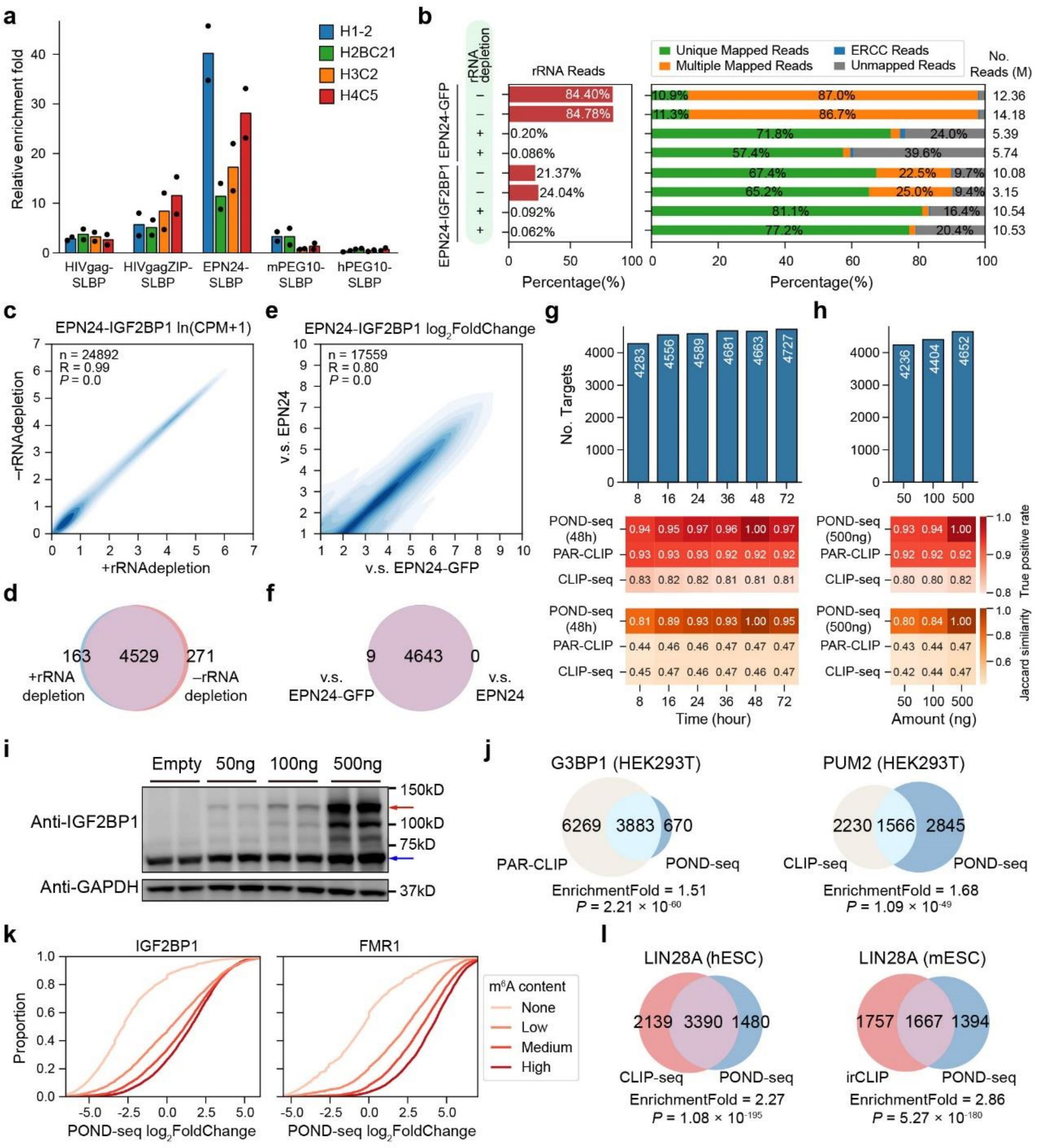

**Supplementary Fig.1: Developing and optimizing POND-seq for profiling RBP targets in living cells, related to Fig.1.**

**a,** Bar plots showing RT-qPCR results for histone genes of various kinds of VLPs and EPN24 fused with SLBP. Data were normalized to a spike-in RNA and then to corresponding GFP fusion constructs. Shown are mean, $n = 2$ biological replicates.

**b,** Bar plots showing percentage of rRNA reads (left) and the complexity of POND-seq library reads (right), with and without rRNA depletion.

**c,** Pearson's correlation of ln(CPM+1) value of RNA transcripts between IGF2BP1-POND-seq datasets with and without rRNA depletion.

**d,** Venn diagram showing the overlap of IGF2BP1 targets identified in samples with and without rRNA depletion.

**e,** Pearson's correlation analysis of log2foldchange CPM of RNA transcripts in EPN24-IGF2BP1 versus EPN24-GFP and EPN24 POND-seq.

**f,** Venn diagram showing the overlap of IGF2BP1 targets identified in EPN24-IGF2BP1 versus EPN24-GFP and EPN24 POND-seq.

**g,** Bar plots showing the number of IGF2BP1 targets identified in POND-seq conducted over varying durations of nanocage production. Heatmaps display the true positive rate (top, red) and Jaccard similarity coefficient (bottom, orange) comparing IGF2BP1 targets identified by POND-seq with those identified by other methods.

**h,** Bar plots showing the number of IGF2BP1 targets identified by POND-seq with varying dosages of nanocage plasmid transfection. Heatmaps display the true positive rate (top, red) and Jaccard similarity coefficient (bottom, orange) comparing IGF2BP1 targets identified by POND-seq with those identified by other methods.

**i,** Western blot showing EPN24-IGF2BP1 protein levels in HEK293T cell transfected with varying dosages of nanocage plasmid. The red arrow refers to EPN24-IGF2BP1, and blue arrow refers to endogenous IGF2BP1. GAPDH was used as the sample loading control.

**j,** Venn diagrams showing the overlap of G3BP1 (left) and PUM2 (right) targets identified by POND-seq and other methods in HEK293T cells.

**k,** Cumulative plots showing the distribution of $\log_2$foldchange for transcripts in EPN24-IGF2BP1 versus EPN24 and EPN24-FMR1 versus EPN24 POND-seq in HEK293T cells. Transcripts were divided into 4 groups based on $m^6A$ content measured by GLORI (None $n$ = 30,987, Low $n$ = 5,085, Medium $n$ = 5,085, High $n$ = 5,085).

**l,** Venn diagrams showing the overlap of LIN28A targets identified by POND-seq and other methods in hESC (left) and mESC (right) cells.

The correlation coefficient R and *P*-values in (**c**) and (**e**) were determined by two-tailed Pearson's correlation test. In (**j**) and (**l**), *P*-values were determined by two-tailed Fisher's exact test.

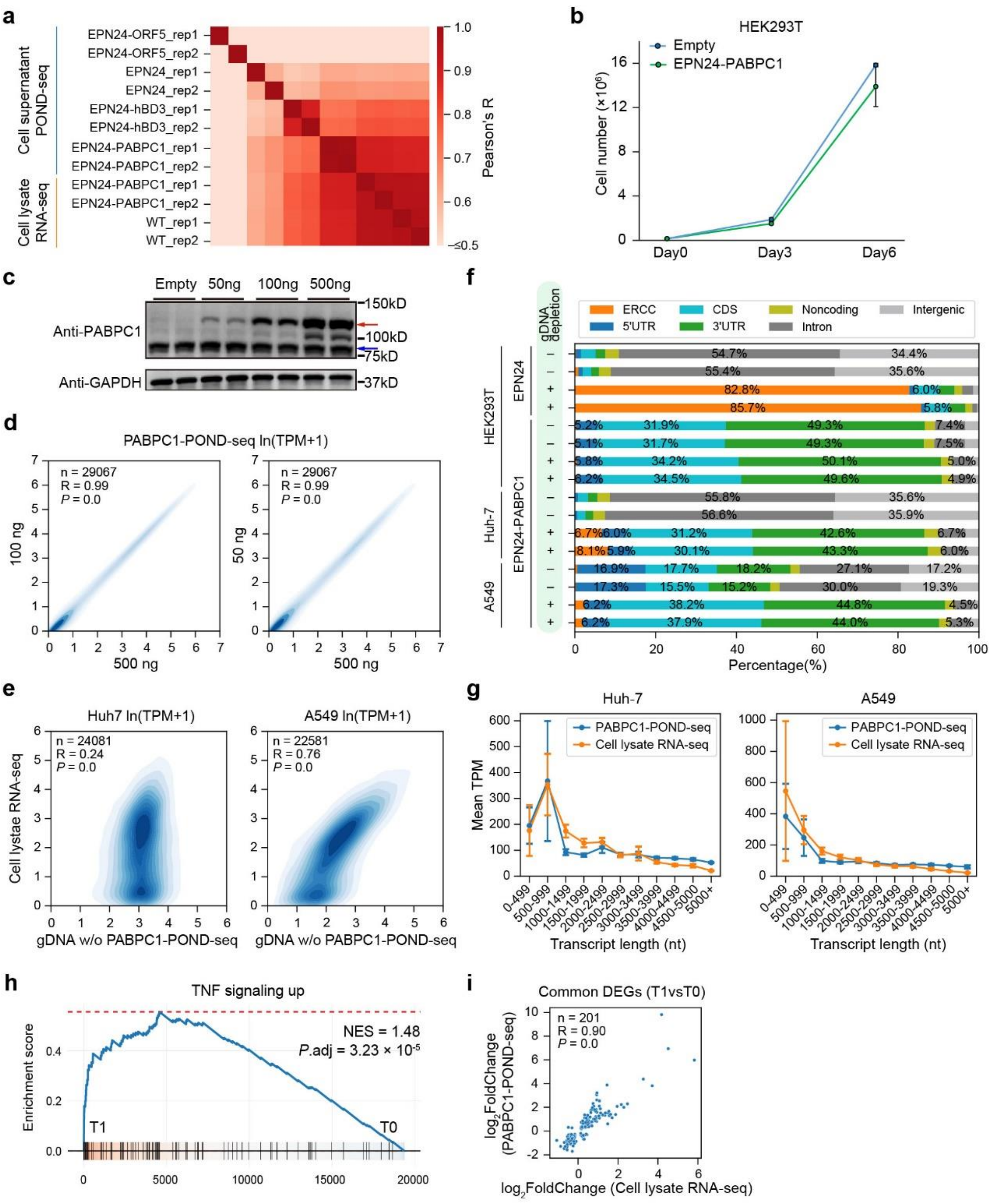


**Supplementary Fig.2: PABPC1-POND-seq captures transcriptome in a non-destructive manner, related to Fig.2.**

**a,** Heatmap showing the correlation of transcriptomes determined by POND-seq with various nanocage constructs and cell lysate RNA-seq. The color intensity indicates Pearson's correlation coefficient.

**b,** Line plot showing proliferation of HEK293T cells transfected with EPN24-PABPC1 or empty plasmid. Shown are mean ± SD, $n = 3$ independent experiments.

**c,** Western blot displaying EPN24-PABPC1 protein levels in HEK293T cells transfected with varying dosages of EPN24-PABPC1 plasmid. The red arrow refers to EPN24-PABPC1, and blue arrow refers to endogenous PABPC1. GAPDH was used as the sample loading control.

**d,** Pearson's correlation of transcriptomes by PABPC1-POND-seq conducted with varying dosages of EPN24-PABPC1 plasmid in HEK293T cells.

**e,** Pearson's correlation of transcriptomes determined by cell lysate RNA-seq and PABPC1-POND-seq without removal of gDNA in Huh-7 and A547 cells.

**f,** Bar plots showing the complexity of reads in PABPC1-POND-seq with or without gDNA removal in HEK293T, Huh-7, and A547 cells.

**g,** Line plots illustrating the mean transcripts per million (TPM) of expressed coding transcripts (TPM >1) in Huh-7 (left) and A549 (right) cells across bins of transcript lengths. Transcriptome data determined by PABPC1-POND-seq are compared to those determined by the corresponding cell lysate RNA-seq. Each data point represents the mean ± SD TPM value for transcripts within the corresponding length bin. Shown are averages of two biological replicates.

**h,** Enrichment profiles showing that TNF signaling upregulated genes were significantly increased following TNFα treatment as determined by PABPC1-POND-seq.

**i,** Scatter plots illustrating the correlation of log2foldchange in TPM for common differentially expressed genes (DEGs) upon TNFα treatment between PABPC1-POND-seq and cell lysate RNA-seq in HEK293T cells.

The correlation coefficient R and *P*-values in (**d**), (**e**), and (**i**) were determined by two-tailed Pearson's correlation test.

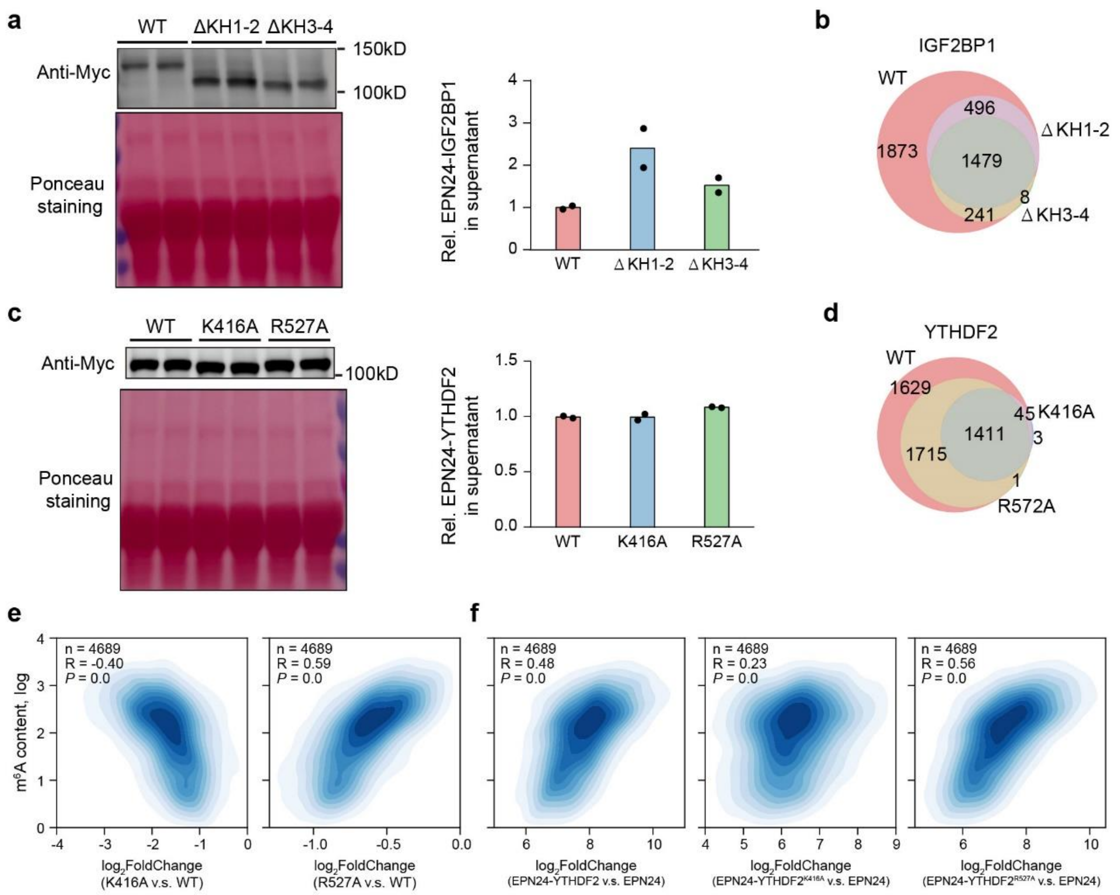


**Supplementary Fig.3: Functional dissection of RBP domains and residues by POND-seq, related to Fig.3.**

**a,** Western blot (top) showing EPN24-IGF2BP1, EPN24-IGF2BP1$^{\Delta KH1\text{-}2}$, and EPN24-IGF2BP1$^{\Delta KH3\text{-}4}$ protein level in supernatant. Ponceau S staining was used to assess the sample loading amount (bottom). Bar plots (right) showing the quantification of protein levels. Show are mean, $n = 2$ biological replicates.

**b,** Venn diagrams showing the overlap of wild type IGF2BP1, IGF2BP1$^{\Delta KH1\text{-}2}$, and IGF2BP1$^{\Delta KH3\text{-}4}$ targets identified by POND-seq.

**c,** Western blot (top) showing wild type YTHDF2, YTHDF2$^{K416A}$, and YTHDF2$^{R527A}$ protein level in supernatant. Ponceau S staining was used to assess the sample loading amount (bottom). Bar plots (right) showing the quantification of protein levels. Show are mean, $n = 2$ biological replicates.

**d,** Venn diagrams showing the overlap of wild type YTHDF2, YTHDF2$^{K416A}$, and YTHDF2$^{R527A}$ targets identified by POND-seq.

**e,** Correlation of $m^6A$ content and $log_2$foldchange of TPM for YTHDF2 bound transcripts in EPN24-YTHDF2$^{K416A}$ (left) and EPN24-YTHDF2$^{R527A}$ (right) versus EPN24-YTHDF2 (WT) POND-seq in HEK293T cells.

**f,** Correlation of $m^6A$ content and $log_2$foldchange of TPM for YTHDF2 bound transcripts in EPN24-YTHDF2 (WT, left), EPN24-YTHDF2$^{K416A}$ (middle) and EPN24-YTHDF2$^{R527A}$ (right) versus EPN24 (control) POND-seq in HEK293T cells.

In (**e**) and (**f**), $m^6A$ content was measured by GLORI, and the correlation coefficient R and *P*-values were determined by two-tailed Pearson's correlation test.

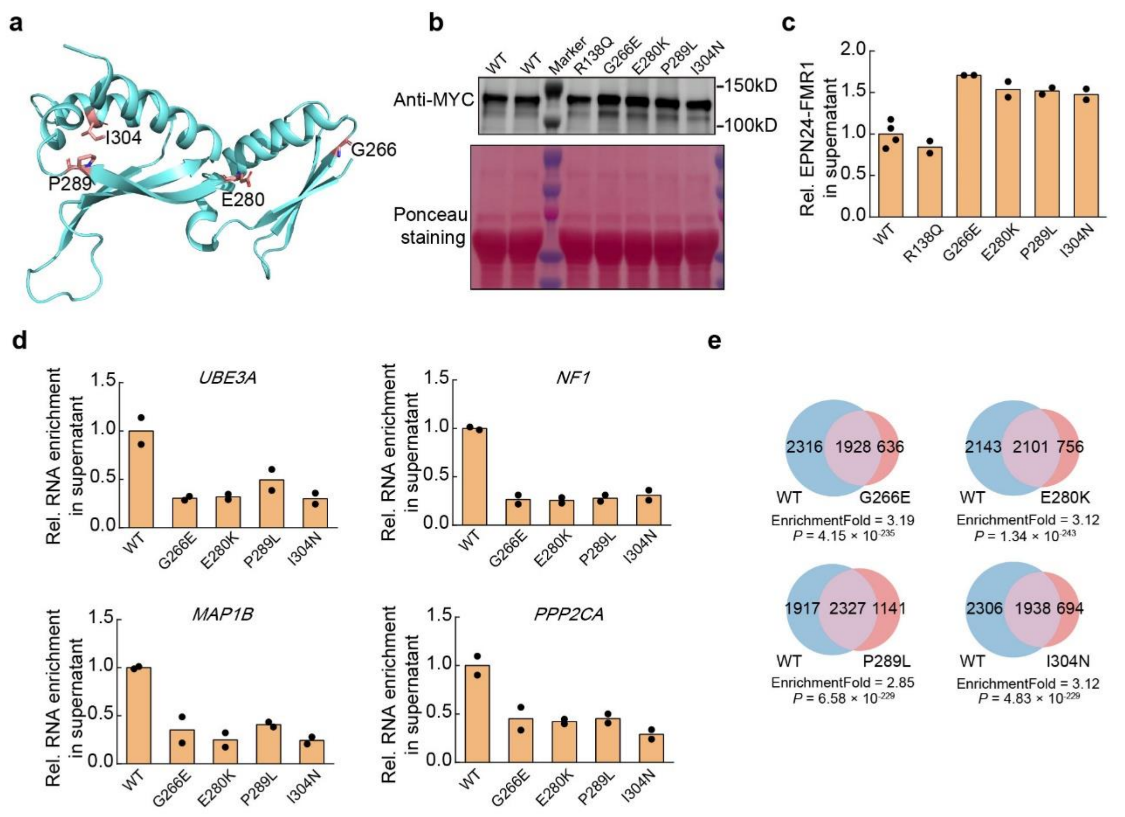


**Supplementary Fig.4: Large scale POND-seq screening identifies new pathogenic mutations in FMR1, related to Fig.4.**

**a,** Location of G266, E280, P289, and I304 in crystal structure of the KH1-KH2 domain of FMR1 (RMSD 0.612 Å for 115 aligned Cα atoms).

**b,** Representative image of western blot showing protein levels of wild-type and various mutant EPN24-FMR1 in the supernatant. Ponceau S staining was used to assess the sample loading amount.

**c,** Bar plots displaying the quantification of protein levels in (**b**). Data were normalized to wild type EPN24-FMR1 level. Show are mean, $n$ = 2 biological replicates.

**d,** RT-qPCR detection of target RNAs bound to WT and mutant FMR1 in the supernatant. Data were normalized to a spike in RNA and then to WT FMR1.Shown are mean, $n$ = 2 biological replicates.

**e,** Venn diagrams showing the overlap of target RNAs bound to G266E, E280K, P289L, and I304N variants compared to wild type FMR1, as determined by POND-seq. *P*-values were determined by two-tailed Fisher's exact test.
